# Microfilaremic loiasis is associated with T cell hyporesponsiveness against SARS-CoV-2

**DOI:** 10.64898/2026.03.02.708985

**Authors:** Maximilian Augé Stock, Dearie Glory Okwu, Ayong More, Anna Doralt, Rodrigue Bikangui, Ismaël Pierrick Mikelet Boussoukou, Kirsten Alexandra Eberhardt, Maren Sandkuhl, Rella Zoleko Manego, Ghyslain Mombo-Ngoma, Matthew McCall, Minka Breloer, Meral Esen, Marylyn Addo, Bertrand Lell, Luzia Veletzky, Rafiou Adamou, Maria Sophia Mackroth

**Author notes:** corresponding authors (LV), (MSM). These authors contributed equally to this work (Luzia Veletzky, Rafiou Adamou, Maria Sophia Mackroth).

## Abstract

**Background:** Loiasis is a chronic filarial infection endemic to Central and West Africa. Although long considered benign, increasing evidence links loiasis to substantial morbidity and mortality. The infection is associated with immune modulation, including Th2-skewed responses and elevated regulatory cytokines. Clinically, loiasis is classified as microfilaremic (presence of circulating microfilariae) or amicrofilaremic (“occult”) disease, the latter defined by a history of eyeworm migration without detectable microfilaremia. This study investigated how chronic *L. loa* infection influences antibody and T cell responses to SARS-CoV-2 following natural infection.

**Methods:** Between 2022 and 2024 this cross-sectional study was done in Lambaréné and surrounding rural areas of Gabon. Study procedures included diagnostics for loiasis and immunological assays. Microfilaremia was confirmed by stained blood smear microscopy, and occult disease was identified using the Rapid Assessment Procedure for Loiasis. SARS-CoV-2-specific IgG responses to spike and nucleocapsid proteins were measured by ELISA, and IFN-γ responses to spike antigen were assessed using an interferon-gamma release assay.

**Results:** Overall, 192 participants were categorized as microfilaremic (n=43), occult loiasis (n=59), or without evidence of active loiasis (n=90). IFN-γ responses were reduced in microfilaremic individuals compared with other participants (*p*= 0.031), whereas IgG responses did not differ. Subsequent analysis across the three groups confirmed that IFN-γ responses were lower in microfilaremic compared with occult participants (*p*= 0.012).

**Conclusion:** These findings suggest that microfilaremic loiasis may impair proinflammatory T cell responses to viral antigens, highlighting the need for further research into the broader immunological effects of *Loa loa* infection in endemic populations.

**Author’s summary:** Loiasis is a parasitic infection caused by the worm *Loa loa* and is common in parts of Central and West Africa. Although long considered relatively benign, growing evidence indicates that loiasis is associated with substantial morbidity. The immunological consequences of chronic *Loa loa* infection remain poorly understood. A small number of studies suggest that *Loa loa* may influence immune regulation, but its broader impact on antiviral immunity is largely unknown.

The COVID-19 pandemic provided a unique opportunity to examine immune responses to a newly emerging virus in a population where loiasis is endemic. We therefore investigated how different forms of *Loa loa* infection influence immune responses after natural SARS-CoV-2 infection. We compared individuals with circulating microfilariae in their blood (microfilaremic), individuals with occult loiasis (history of eye worm), and individuals without signs of active infection.

We found that microfilaremic individuals had weaker virus-specific IFN-γ T cell responses, while antibody levels were similar across groups. These findings suggest that active loiasis may dampen certain antiviral immune functions. Understanding the underlying mechanisms is important, as such immune modulation could affect responses to vaccines and other infectious diseases in endemic regions.

## Introduction

*Loa loa*, commonly known as the “African eyeworm”, is a filarial parasite endemic to Central and West Africa. Transmission occurs through the bites of *Chrysops* flies and recent estimates consider millions of individuals to be infected [1]. Historically considered benign, *L. loa* infection, or loiasis, has received limited public health attention and is notably absent from the WHO’s list of neglected tropical diseases (NTDs). However, recent evidence links loiasis to significant morbidity and increased mortality, prompting a re-evaluation of its clinical and public health impact [2,3]. Infected individuals may present with microfilaremia, defined by detectable circulating *L. loa* microfilariae in the blood, or as amicrofilaremic (occult) loiasis, in which clinical symptoms occur without detectable microfilariae, most notably adult worm migration across the eye [4]. The determinants of these divergent clinical presentations are not fully understood but likely involve a combination of host genetic and immunological as well as parasite factors [4–6].

Helminth infections, including loiasis, are known to modulate host immunity, often shifting responses toward a Th2-type profile and enhancing regulatory pathways. This includes elevated IL-10 and TGF-β production and regulatory T cell activation, which can suppress effector T cell responses to various antigens [7–14]. Such modulation may impair responses to both vaccines and unrelated pathogens. Although data specific to *L. loa* are limited, existing studies show distinct immunological profiles between microfilaremic and amicrofilaremic individuals. Amicrofilaremic individuals have been shown to exhibit stronger Th1 and Th2 responses as well as increased T cell proliferation in response to *L. loa* antigen, whereas microfilaremic individuals showed marked T cell hyporesponsiveness [15]. Additionally, higher levels of IL-4, IL-5, and filarial-specific antibodies in amicrofilaremic patients compared to their microfilaremic counterparts have been described; these differences disappeared one year after treatment [16]. Conversely, a third study observed that microfilaremic individuals mounted stronger Th2 and IL-10 responses to mitogens compared to individuals with no signs of active infection [17].

These immune differences may affect responses to co-infections, as evidenced by a study reporting that HIV-infected microfilaremic individuals exhibited lower CD4^+^ T cell counts and higher viral loads compared with amicrofilaremic individuals [18]. Despite such findings, the broader implications of *L. loa*-induced immunomodulation remain poorly understood. The emergence of the COVID-19 pandemic provided a unique opportunity to examine how chronic loiasis may affect host immune responses to a new viral infection. To this end, this study investigated the impact of microfilaremic and amicrofilaremic *L. loa* infection on immune responses to SARS-CoV-2 in unvaccinated individuals. We hypothesized that chronic loiasis impairs SARS-CoV-2-specific immune responses.

## Material and methods

### Ethics statement

The study was performed in compliance with the International Conference for Harmonization (ICH) - Good Clinical Practices (GCP) - E6 (R2) and local ethical requirements. Ethical approval was obtained from the Comité d’Ethique Institutionnel du Centre de Recherches Médicales de Lambaréné (CEI-009/2022) and the Ethics Committee of the Medical Association of Hamburg (101501-BO-ff). Written informed consent was taken from all study participants before any study related procedures were initiated.

### Study design and participants

This cross-sectional study was conducted between 2022 and 2024 in rural villages of the Ngounié and Moyen-Ogooué provinces of Gabon. Eligible participants were adults (≥18 years) who had not received SARS-CoV-2 vaccination and were not pregnant. All individuals who expressed interest and provided written informed consent were enrolled. The participants received detailed information about the study procedures prior to enrollment. Additionally, antibody assays were also performed in 30 samples from a study conducted in Gabon before the SARS-CoV-2 pandemic to assess baseline cross reactivity in the study population [2].

### Sample and data collection

Participants completed a standardized questionnaire that captured demographic information and the Rapid Assessment Procedure for Loiasis (RAPLOA), for history of eyeworm [19]. Venous blood (11 mL) was collected between 9:30 AM and 4:30 PM on the day of recruitment to account for the diurnal periodicity of *L. loa* microfilaremia. Hematological analyses, full blood count and leukocyte differentiation, were performed on the day of inclusion.

Stool and urine samples were either collected at the time of recruitment or retrieved later using pre-distributed containers. All samples were processed at the Centre de Recherches Médicales de Lambaréné (CERMEL), Gabon.

### Parasitological examinations

At least 40µL of venous blood were prepared as Giemsa-stained thick blood smears, following established protocols [2]. Blood smears were examined independently by two microscopists for detection of *L. loa* and *Mansonella* spp. microfilaria. If results differed by more than 50%, a third reading was conducted. Microfilaria counts were extrapolated to 1 mL of blood. Additionally, thick blood smears were examined for detection of *Plasmodium* spp. Stool and urine samples were examined to identify soil-transmitted helminths, intestinal protozoa and Schistosoma eggs using established protocols [20].

### Interferon-γ T-cell responses

Heparinized blood was processed according to the Quan-T-Cell SARS-CoV-2 kit (Euroimmun:ET2606-3003) protocol to stimulate T cells. Samples were incubated for 22 hours, centrifuged and the supernatants analyzed for Interferon-gamma (IFN-γ) levels using the Quan-T-Cell ELISA (Euroimmun: EQ6841-9601), following the manufacturer’s instructions. The remaining heparinized blood was centrifuged at 1000 RCF for 15 minutes to obtain plasma for antibody analyses. IFN-γ concentrations were validated according to kit instructions. Invalid results were excluded from analysis. Samples with high blank values (>400 mIU/mL) were retained if the positive control exceeded 200% of the blank, accounting for higher background IFN-γ levels in this cohort. A stimulation response was considered negative if it did not exceed 150% of the blank value. The relative validation criteria for the IFN-γ release assay led to some Quan-T-Cell-test-negative individuals having a higher measured IFN-γ concentration than some positive individuals. Therefore, negative individuals were excluded from subsequent statistical analysis.

### Antibody responses

Plasma was analyzed at CERMEL using the Anti-SARS-CoV-2-QuantiVac-ELISA (IgG)-Kit (Euroimmun: EI2606-9601-10G) to quantify antibodies targeting the spike protein S1. Analyses followed the manufacturer’s recommendations, with minor changes to the washing protocol. Additional plasma aliquots were transferred to Hamburg, Germany, to measure the nucleocapsid-specific IgG using the Anti-SARS-CoV-2-NCP (IgG) ELISA-Kit (Euroimmun:2606-9601-2G) according to manufacturer’s recommendations.

### Data management

All personal identifiers were pseudonymized with unique participant codes, accessible only to key research personnel. Trained staff manually entered and double-checked data for accuracy. Physical records were stored in locked facilities at CERMEL, and electronic data were managed in REDCap with regular institutional backups

### Statistical analysis

Participants in Gabon were categorized based on *L. loa* diagnostics: 1. No sign of active loiasis: No detectable microfilaremia and a negative eyeworm history (RAPLOA), 2. Occult loiasis: No detectable microfilaremia but a positive eyeworm history, 3. Microfilaremic loiasis: Detectable microfilaremia, regardless of eyeworm history. For primary analyses, all non-microfilaremic individuals, comprising individuals with occult infection as well as individuals with no sign of active loiasis, were compared to microfilaremic participants.

Continuous variables were summarized as medians and interquartile ranges (IQR) and compared using Wilcoxon rank sum or Kruskal–Wallis tests. For variables showing statistical significance, pairwise post hoc tests were conducted with false discovery rate (FDR) correction for multiple comparisons. Additionally, multiple linear regression models were fitted to assess group differences adjusted for age and sex as potential confounders. Dependent variables were normalized using the R package bestNormalize prior to analysis. All tests were two-sided with significance at p < 0.05. Statistical analyses were performed in R (version 4.4.3, R Foundation for Statistical Computing, Vienna, Austria). The study is reported aligning with the STROBE statement.

## Results

### Study Population

A total of 193 Loa-exposed individuals, comprising 108 women and 85 men, aged 19–94 years, were enrolled in the study. One male participant was excluded from statistical analysis due to missing RAPLOA assessment. Demographic details, loiasis states, and parasitological coinfections are summarized in table 1.

**Table 1.** Overview of the study population, by demographic data, loiasis infection state and parasitic coinfections.

|  | Overall<br>(N=192<sup>±</sup>,<br>column %) | No sign of<br>loiasis<br>(n=90,<br>column %) | Occult<br>loiasis<br>(n=59,<br>column %) | Microfilaremic<br>loiasis<br>(n=43,<br>column %) |
| --- | --- | --- | --- | --- |
| **Sex** |  |  |  |  |
| Female | 108 (56.2%) | 56 (62.2%) | 36 (61%) | 16 (37.2%) |
| Male | 84 (43.8%) | 34 (37.8%) | 23 (39%) | 27 (62.8%) |
| **Age group<sup>§</sup>** |  |  |  |  |
| 19-44 years | 64 (33.5%) | 30 (33.7%) | 20 (33.9%) | 14 (32.6%) |
| 45-69 years | 100 (52.4%) | 45 (50.6%) | 31 (52.5%) | 24 (55.8%) |
| 70-94 years | 27 (14.1%) | 14 (15.7%) | 8 (13.6%) | 5 (11.6%) |
| **Region** |  |  |  |  |
| Rural | 70 (36.5%) | 29 (32.2%) | 26 (44.1%) | 15 (34.9%) |
| Semi-urban | 122 (63.5%) | 61 (67.8%) | 33 (55.9%) | 28 (65.1%) |
| ***Loa loa* microfilaremia per mL (median, IQR)** |  | 0 | 0 | 990 (181-3698) |
| **Parasitic coinfection** |  |  |  |  |
| Coinfection unknown* | 31 (16.1%) | 13 (14.4%) | 9 (15.3%) | 9 (20.9%) |
| No coinfection | 67 (34.9%) | 31 (34.4%) | 23 (39.0%) | 13 (30.2%) |
| Any coinfection | 94 (49.0%) | 46 (51.1%) | 27 (45.8%) | 21 (48.8%) |
| *Plasmodium* spp. | 30 (15.6%) | 14 (15.6%) | 10 (16.9%) | 6 (14%) |
| *Mansonella* spp. | 11 (5.7%) | 3 (3.3%) | 3 (5.1%) | 5 (11.6%) |
| *Schistosoma* spp. | 14 (7.3%) | 6 (6.7%) | 6 (10.2%) | 2 (4.7%) |
| STH spp.** | 17 (8.9%) | 8 (8.9%) | 2 (3.4%) | 7 (16.3%) |
| Intestinal protozoan spp.<sup>°</sup> | 55 (28.7%) | 30 (33.3%) | 15 (25.4%) | 10 (23.3%) |

| Overall<br>(N=192<sup>‡</sup>,<br>column %) | No sign of<br>loiasis<br>(n=90,<br>column %) | Occult<br>loiasis<br>(n=59,<br>column %) | Microfilaremic<br>loiasis<br>(n=43,<br>column %) |
| --- | --- | --- | --- |
| <sup>‡</sup>data of the one excluded individual are not shown. <sup>§</sup>for 1 individual age data missing <sup>*</sup>participants that didn't return a full set of blood, urine and stool sample. <sup>**</sup>Soil-transmitted-Helminths: *Trichuris trichiura*, *Ascaris lumbricoides*, *Necator americanus*, *Strongyloides stercoralis*. <sup>°</sup>Intestinal Protozoa: *Giardia lamblia*, *Entamoeba histolytica*, *Entamoeba coli*, *Cyclospora cayetanensis*, *Iodaemoeba butschlii*, *Endolimax nana*, *Blastocystis hominis*, *Chilomastix mesnili*; and *Mansonella* spp.. <sup>**</sup> |  |  |  |

### Hematological findings

Hemoglobin levels did not differ between the loiasis groups (Fig 1). In contrast, *L. loa* microfilaremic individuals had significantly higher leukocyte counts compared to individuals with occult loiasis and to those with no sign of loiasis (6.4 vs. 5.0 G/L for each comparison, *p*=0.001). Microfilaremic individuals also had higher absolute and relative eosinophil counts (0.9 G/L and 19% vs. 0.3 G/L and 7%, and vs. 0.3 G/L and 6%, respectively; *p*=0.001 for all comparisons) than the other two groups. These differences remained significant after adjustment for age and sex (table 2).

**Table 2.** Overview on outcomes of hematological and immunological analyses by groups as well as group comparisons.

|  | Overall<br>Median<br>(IQR) | NL<br>Median<br>(IQR) | OL<br>Median<br>(IQR) | ML<br>Median<br>(IQR) | p-<br>value<sup>°</sup> | NL vs.<br>OL<sup>#</sup> | Adj. NL<br>vs. OL<sup>*</sup> | ML vs.<br>NL<sup>#</sup> | Adj. ML<br>vs. NL<sup>*</sup> | ML vs.<br>OL<sup>#</sup> | Adj. ML<br>vs. OL<sup>*</sup> |
| --- | --- | --- | --- | --- | --- | --- | --- | --- | --- | --- | --- |
| **Hemoglobin (g/dL)** | 12<br>(11-13)<br>[n = 175] | 12<br>(11-13)<br>[n = 82] | 12<br>(11-13)<br>[n = 54] | 12<br>(11-13)<br>[n = 39] | 0.494 | NA | NA | NA | NA | NA | NA |
| **Leucocytes (G/L)** | 5.2<br>(4.2-6.5)<br>[n = 175] | 5.0<br>(4.2-6.0)<br>[n = 82] | 5.0<br>(3.8-6.3)<br>[n = 54] | 6.4<br>(5.2-7.2)<br>[n = 39] | **0.001** | 0.784 | 0.672 | **0.001** | **0.006** | **0.001** | **0.004** |
| **Eosinophils (rel.%)** | 10<br>(3-20)<br>[n = 163] | 6<br>(3-16)<br>[n = 77] | 7<br>(2-21)<br>[n = 49] | 19<br>(13-26)<br>[n = 37] | **0.001** | 0.770 | 0.833 | **0.001** | **<0.001** | **0.001** | **0.002** |
| **Eosinophils (G/L)** | 0.5<br>(0.1-1.1)<br>[n = 163] | 0.3<br>(0.1-0.8)<br>[n = 77] | 0.3<br>(0.1-1.0)<br>[n = 49] | 0.9<br>(0.7-1.7)<br>[n = 37] | **0.001** | 0.990 | 0.879 | **0.001** | **<0.001** | **0.001** | **<0.001** |
| **Anti-S1-IgG (BAU/mL)** | 200<br>(101-359)<br>[n = 192] | 190<br>(107-357)<br>[n = 90] | 240<br>(80-446)<br>[n = 59] | 180<br>(102-302)<br>[n = 43] | 0.695 | NA | NA | NA | NA | NA | NA |
| **Anti-NCP-IgG (Ratio)** | 0.8<br>(0.5-1.6)<br>[n = 192] | 0.8<br>(0.5-1.6)<br>[n = 90] | 0.9<br>(0.5-1.9)<br>[n = 59] | 0.7<br>(0.4-1.4)<br>[n = 43] | 0.474 | NA | NA | NA | NA | NA | NA |
| **Interferon-γ**<br>**concentration (mIU/mL)** | 1177<br>(684-2406)<br>[n = 162] | 1451<br>(634-2300)<br>[n = 75] | 1479<br>(833-3031)<br>[n = 51] | 870<br>(585-1353)<br>[n = 36] | **0.019** | 0.257 | 0.191 | 0.078 | 0.118 | **0.015** | **0.012** |

IQR – Interquartile range. <sup>°</sup> crude p-value from global Kruskal-Wallis test, <sup>#</sup> FDR-corrected pairwise p-values from post hoc tests following Kruskal-Wallis test, <sup>\*</sup>p-value adjusted for sex and age from multiple linear regression, NL = No sign of active loiasis, OL= Occult loiasis, ML = Microfilaremic loiasis.

**Fig 1.**
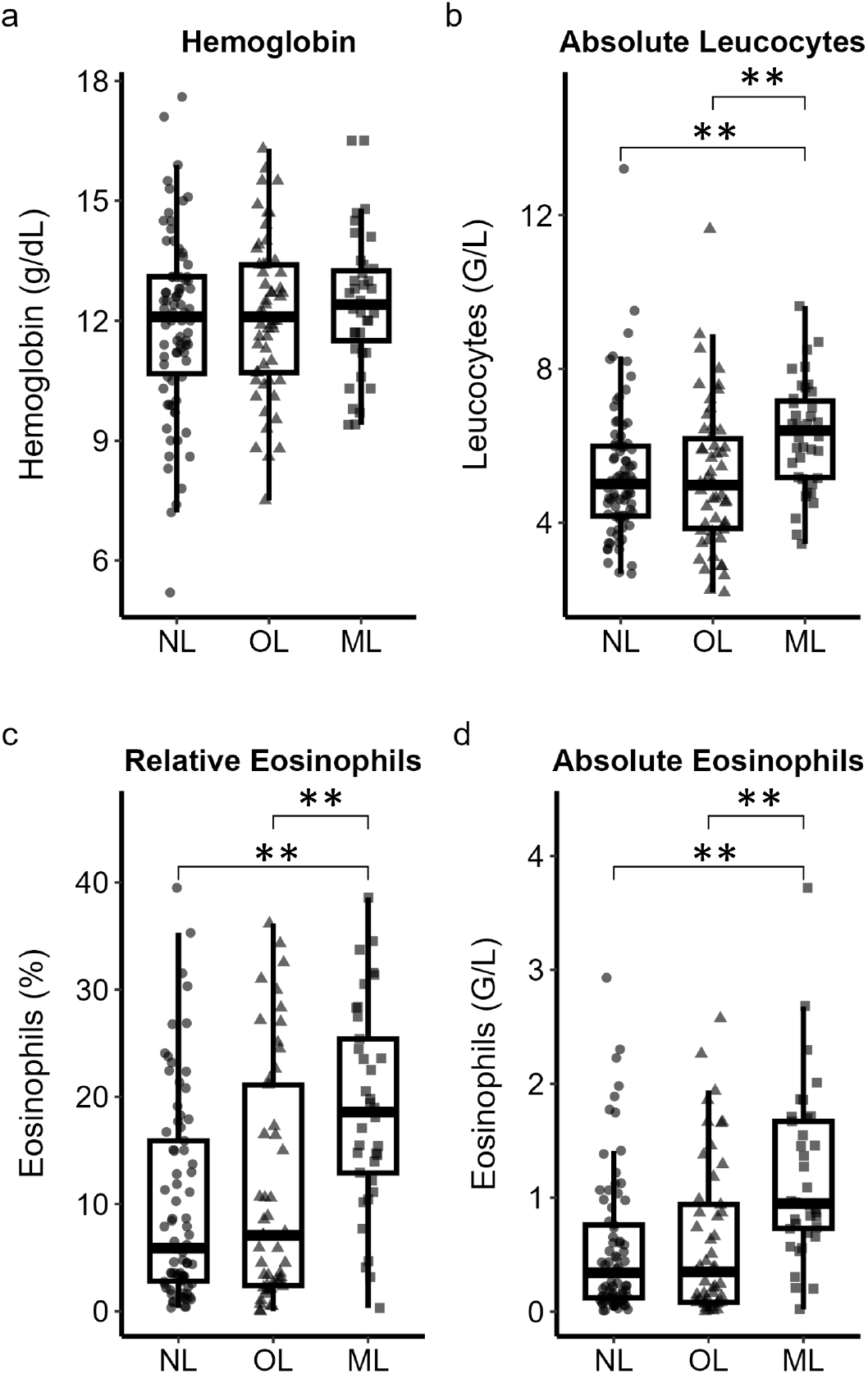
Overview of hematological findings. Results are shown as individual values and boxplots for hemoglobin (a), leukocyte count (b), relative (c), and absolute eosinophil count (d). The box shows the median and interquartile range (IQR, 25th–75th percentile). Whiskers extend to the most extreme values not exceeding 1.5 × IQR from the first and third quartiles; outliers beyond these limits are plotted as individual points. Intergroup differences were assessed using non-parametric Kruskal Wallis and corrected for multiple testing using the false discovery method (FDR). *P*-values of intergroup comparisons are provided if *p*<0.05. \*\**p*< 0.01. NL=No sign of active loiasis, OL=Occult loiasis, ML=Microfilaremic loiasis.

### Immune responses to SARS-CoV-2 in Loa-exposed individuals

Anti-S1-IgG and anti-NCP-IgG results were available for all possibly loiasis exposed participants (n=192), see table 2. Among those, 14 participants (7.3%) were anti-S1-IgG negative, comprising 5 with no sign of active loiasis, 7 with occult loiasis and 2 with microfilaremic loiasis, respectively. Further 93 individuals (48.4%), were anti-NCP-IgG negative, comprising 42 with no sign of active loiasis, 26 with occult loiasis and 25 with microfilaremic loiasis. Test negativity was not associated with sex, age, or loiasis state.

IFN-γ T-cell response results were not available for three participants, due to insufficient blood sample amount (1.6%). Further 21 participants (10.9%) were excluded from statistical analysis due to negative responses to SARS-CoV-2 and 6 participants due to invalid results. Hence, results were available for 162 Loa-exposed participants (84.4%). To test for unspecific cross reactions in the antibody assays, 30 prepandemic samples from the same study area were analyzed for the presence of anti-S1 and anti-NCP antibodies. Of these samples, three had a weakly positive S1 antibody and three a borderline positive NCP result (S1 Fig).

First, results were compared between microfilaremic versus non-microfilaremic individuals, which comprised the groups of no sign of active infection and occult infection (Fig 1 upper panel and S1 table). No differences were observed between microfilaremic and non-microfilaremic in the tested SARS-CoV-2-specific antibody responses. However, IFN-γ concentrations were lower in microfilaremic compared with non-microfilaremic participants (median (IQR) mIU/mL: microfilaremic 870 (585-1353) vs. non-microfilaremic 1461 (733-2844), *p*=0.01). This difference remained significant after adjustment for confounders (*p*= 0.031).

Subsequent comparisons across loiasis subgroups showed no statistically significant differences in anti-S1-IgG concentrations and anti-NCP-IgG ratios between the groups. However, measured IFN-γ concentrations were lower in microfilaremic individuals compared to those with occult loiasis (median (IQR) mIU/mL: microfilaremic 870 (585-1353) vs. occult 1479 (833-3031), *p*= 0.015, see Fig 2, lower panel). The observed difference between microfilaremic and occult remained significant after adjustment for confounders (*p*=0.012). Absolute IFN-γ and absolute eosinophil numbers showed a weak negative correlation (rho = - 0.17, *p*=0.045) as did absolute IFN-γ and microfilaremia (rho=-0.2, *p*=0.012), as tested using spearman’s rank correlation coefficient (S2 Fig).

**Fig 2.**
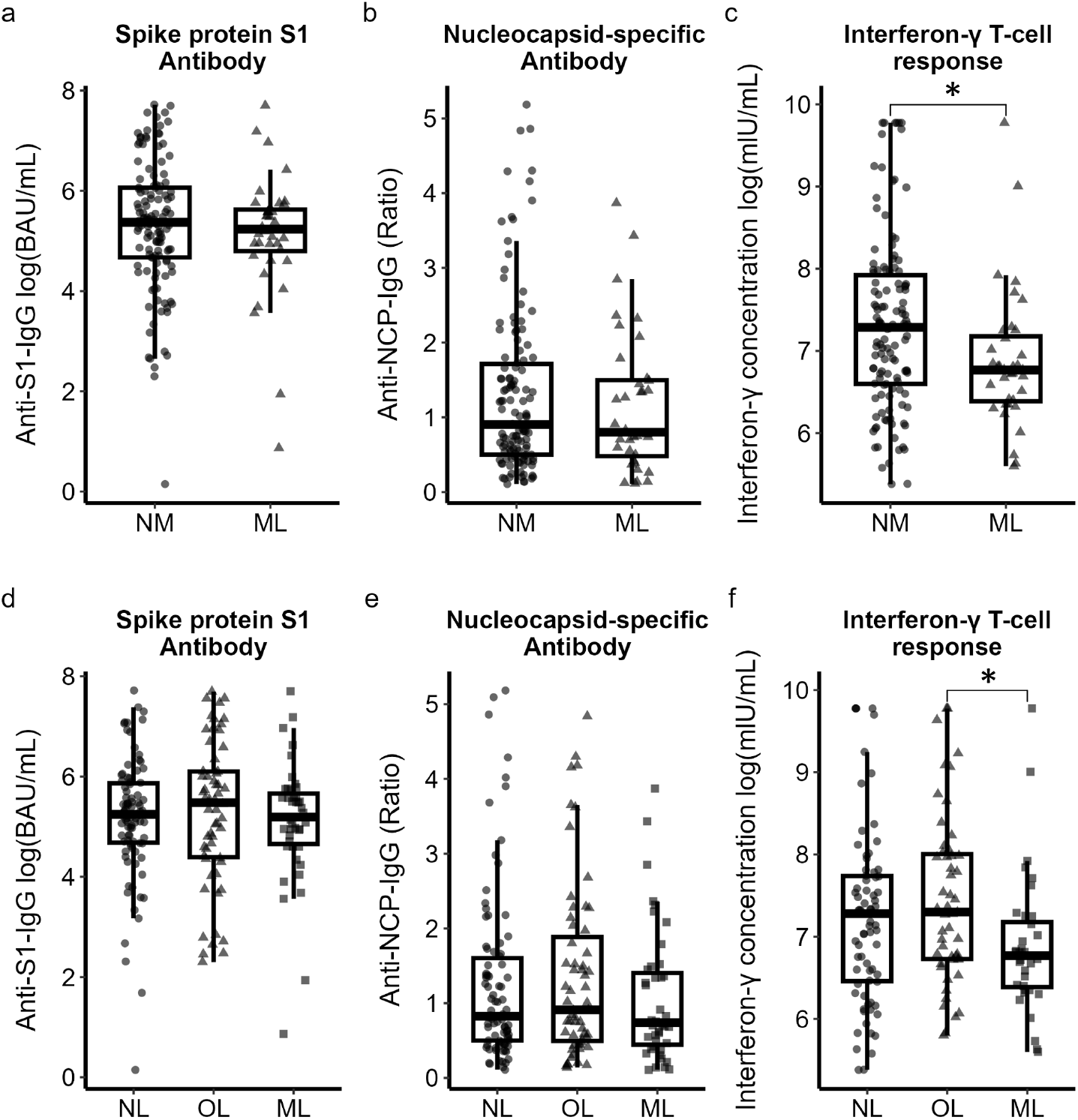
Immune responses to SARS-CoV-2 in loiasis exposed individuals. Upper panel: comparison of non-microfilaremic (= NM: No sign of active loiasis combined with occult loiasis) versus microfilaremic participants (ML = Microfilaremic loiasis). Lower panel: comparison across three groups (NL = No sign of active loiasis, OL= Occult loiasis, ML = Microfilaremic loiasis). For each panel, Anti-S1-IgG concentrations (a), Anti-NCP-IgG ratios (b), Interferon-γ concentrations (c) are shown as individual values. The box shows the median and interquartile range (IQR, 25th–75th percentile). Whiskers extend to the most extreme values not exceeding 1.5 × IQR from the first and third quartiles; outliers beyond these limits are plotted as individual points. Intergroup differences were assessed using non-parametric Wilcoxon-rank-sum test (upper panel) or Kruskal-Wallis with subsequent pairwise post hoc comparisons using false discovery rate (FDR) correction (lower panel). *P*-values of intergroup comparisons are provided if p < 0.05. * p < 0.05.

## Discussion

The immunomodulatory effects of chronic parasitic infections, including reduced immune responses to unrelated, heterologous antigens, are well established. As loiasis is a chronic or even lifelong parasitic infection, it may have a significant impact on the immune system of infected individuals. Therefore, this study aimed to explore whether chronic loiasis alters immune responses to the co-infection SARS-CoV-2, by measuring virus-specific IFN-γ production and IgG responses to the spike (S1) and nucleocapsid (NCP) proteins in unvaccinated individuals.

We found that *L. loa* microfilaremic individuals exhibited reduced SARS-CoV-2-specific IFN-γ responses compared to Loa-exposed but non-microfilaremic individuals and specifically to those with occult loiasis. However, S1 and NCP IgG values did not differ across individuals from different loiasis subgroups.

These findings suggest that microfilaremic loiasis is associated with a selective impairment in longevity of virus-specific cellular immune responses while preserving virus-specific humoral immunity. Such a dissociation aligns with a T cell–focused immunomodulatory effect, potentially mediated by parasite-induced regulatory pathways which preferentially dampen proinflammatory responses [13].

Previous evidence on the impact of chronic loiasis on immunity to heterologous infections remains limited. Stimulation of PBMCs from microfilaremic loiasis patients from West and Central Africa with *Mycobacterium tuberculosis* antigen resulted in a higher proportion of CD4^+^ IL-4^+^ T cells compared to PBMCs from unexposed North American individuals, suggesting a Th2-biased immune profile. However, no differences were observed in IFN-γ– producing CD4^+^ T cells between the two groups [8]. T cell hypo-responsiveness to both tuberculin PPD and *L. loa* antigens has been observed in populations with high *L. loa* transmission compared to low *L. loa* transmission areas [21]. Further, in HIV-therapy-naïve patients, microfilaremic loiasis was associated with lower CD4^+^ counts and higher viral loads [18]. These findings indicate an immunomodulatory effect of chronic loiasis on co-infections. Parallels can be drawn from studies on other chronic filarial infections. Patients with lymphatic filariasis or onchocerciasis displayed reduced IFN-γ responses to tetanus toxoid, with the latter also showing diminished antibody responses [22,23]. In individuals with lymphatic filariasis receiving SARS-CoV-2 vaccination, antibody levels were preserved, although additional *Ascaris* seropositivity was associated with reduced IgA responses [24].

In the *Litomosoides sigmodontis* mouse model, which mimics human filariasis, chronic infection suppressed influenza-specific antibody responses and protective immunity [13,25]. Notably, while antiparasitic treatment partially restored vaccine responsiveness, some degree of immunosuppression persisted, along with an expanded population of LAG3^+^CD49b^+^ Type 1 regulatory T cels (Tr1 cells). These studies point to a mechanistic link between chronic filarial infections, Tr1 cell induction, and T cell suppression. In humans, elevated IL-10 and Tr1 cell expansion have been documented in lymphatic filariasis and onchocerciasis [14,26]. Taken together, these data suggest that Tr1 cell induction and IL-10–driven suppression could be central to the immune response in chronic filarial infections, offering a possible explanation for the reduced SARS-CoV-2-specific IFN-γ we observed in *L. loa* microfilaremic individuals.

While SARS-CoV-2-specific antibody levels were similar across loiasis subgroups, our study did not assess neutralizing capacity or IgG subclass distribution, which may reveal more nuanced differences in humoral function. Notably, previous work has shown reduced total IgG against *L. loa* antigens in microfilaremic individuals [21,27–29]. However, such parasite-specific alterations do not appear to extend to unrelated antiviral responses in our data.

As described previously, eosinophilia, a hallmark of helminth infection, was significantly elevated in microfilaremic participants [4,30]. Eosinophils are associated with type 2 immune responses, but in our cohort, eosinophilia showed only very weak inverse correlation with SARS-CoV-2-specific IFN-γ concentrations and no correlation with antibody responses. This suggests that in this context eosinophilia rather reflects general helminth-driven immune activation is unlikely to represent a major direct driver of antiviral immune modulation.

While we hypothesized that general parasite exposure would suppress immune responses to SARS-CoV-2, other factors besides *L. loa* infection, including environmental exposures or genetic differences, may also influence the immune response to heterologous antigens. The factors determining the loiasis infection state, i.e. microfilaremic or occult disease presentation are not yet fully understood, but are likely to comprise genetic and immunological factors [4– 6]. Consequently, one may argue that the observed difference may not be loiasis specific but rather reflect the host factors determining the loiasis infection state itself. However, this possibility cannot be addressed within the constraints of the present cross-sectional study design.

Assay specificity is an additional important consideration. Although the Euroimmun ELISAs are highly specific, cross-reactivity with endemic coronaviruses cannot be excluded. However, testing pre-pandemic plasma from the same Gabonese region revealed weakly positive S1 and borderline NCP signals in only ∼10% of samples (S1 Fig), suggesting that cross-reactivity is unlikely to account for the overall high S1 antibodies observed. Nevertheless, prior exposure to other coronaviruses may have primed B and T cells, resulting in stronger SARS-CoV-2 responses even in the absence of detectable pre-existing antibodies. However, if cross reactivity contributed to the observed response, this effect would be expected to be similar across the comparison groups and therefore is unlikely to explain the detected differences.

Further limitations of our study need to be considered. Despite thorough testing, further co-infections cannot be excluded and may have influenced immune responses.

Our study was cross-sectional and primarily descriptive, which limits causal inference regarding the impact of *L. loa* microfilaremia on antiviral immune responses. While we observed reduced SARS-CoV-2-specific IFN-γ production in microfilaremic individuals, the underlying mechanisms remain unclear, and factors such as prior viral exposure could not be fully controlled. Despite these limitations, our findings address an important knowledge gap by suggesting a selective modulation of cellular, but not humoral, immunity in the context of microfilaremic loiasis. Future longitudinal and mechanistic studies, including detailed T cell phenotyping, cytokine profiling, and investigation of regulatory pathways such as Tr1 expansion and IL-10–mediated suppression, will be critical to unravel how *L. loa* microfilaremia modulates host immunity and responses to heterologous infections and vaccinations.

## Acknowledgments

We sincerely thank all study participants for their invaluable contribution to this research. We are grateful to the field workers especially Paul Aimé Kombi for their support in participant recruitment and sample collection. We also acknowledge the support of the staff at CERMEL for facilitating laboratory and administrative logistics.

## Supporting information

**S1 Fig: Immune responses in prepandemic individuals displaying individual values for Anti-S1-Antibody-Concentrations (a) and Anti-NCP-Antibody-Ratios (b).** The upper dotted line represents the positivity threshold and the lower dotted line borderline threshold.

**S1 table: Comparison of immunological assays results between non-microfilaremic and microfilaremic.**

**S2 Fig: Spearman’s rank correlation showing weak correlations of absolute IFN-γ and absolute eosinophil numbers (A) as well as correlation of absolute IFN-γ and microfilaremia (B).**

